# Targeting hypoxic habitats with hypoxia pro-drug evofosfamide in preclinical models of sarcoma

**DOI:** 10.1101/2020.10.06.326934

**Authors:** Bruna V. Jardim-Perassi, Wei Mu, Suning Huang, Michal R. Tomaszewski, Jan Poleszczuk, Mahmoud A. Abdalah, Mikalai M. Budzevich, William Dominguez-Viqueira, Damon R. Reed, Marilyn M. Bui, Joseph O. Johnson, Gary V. Martinez, Robert J. Gillies

**Affiliations:** Department of Cancer Physiology, Moffitt Cancer Center, Tampa, US; Guangxi Medical University Cancer Hospital, Nanning Guangxi, China; Department of Integrated Mathematical Oncology, Moffitt Cancer Center, Tampa, US; Nalecz Institute of Biocybernetics and Biomedical Engineering, Polish Academy of Sciences, Poland; Quantitative Imaging Core, Moffitt Cancer Center, Tampa, Florida; Small Animal Imaging Laboratory, Moffitt Cancer Center, Tampa, Florida; Department of Interdisciplinary Cancer Management, Adolescent and Young Adult Program, Moffitt Cancer Center, Tampa, Florida; Department of Pathology, Moffitt Cancer Center, Tampa, Florida; Analytic Microscopy Core, Moffitt Cancer Center, Tampa, Florida; Department of Imaging Physics, The University of Texas M.D. Anderson Cancer Center

**Author notes:** **Corresponding author: Robert J. Gillies**, Department of Cancer Physiology, Moffitt Cancer Center, 12902 USF Magnolia Drive Tampa, FL 33612. These authors contributed equally to this work (Bruna V. Jardim-Perassi, Wei Mu).

## Abstract

Hypoxic regions (habitats) within tumors are heterogeneously distributed and can be widely variant. Hypoxic habitats are generally pan-therapy resistant. For this reason, hypoxia-activated prodrugs (HAPs) have been developed to target these resistant volumes. The HAP evofosfamide (TH-302) has shown promise in preclinical and early clinical trials of sarcoma. However, in a phase III clinical trial, TH-302 did not improve survival in combination with doxorubicin (dox), most likely due to a lack of patient stratification based on hypoxic status. Herein, our goal was to develop deep-learning (DL) models to identify hypoxic habitats, using multiparametric (mp) MRI and co-registered histology, and to non-invasively monitor response to TH-302 in a patient-derived xenograft (PDX) of rhabdomyosarcoma and a syngeneic model of fibrosarcoma (RIF-1). A DL convolutional neural network showed strong correlations (>0.81) between the true hypoxic portion in histology and the predicted hypoxic portion in multiparametric MRI. TH-302 monotherapy or in combination with Dox delayed tumor growth and increased survival in the hypoxic PDX model (p<0.05), but not in the RIF-1 model, which had lower volume of hypoxic habitats. Control studies showed that RIF-1 resistance was due to hypoxia and not to other causes. Notably, PDX tumors developed resistance to TH-302 under prolonged treatment. In conclusion, response to TH-302 can be attributed to differences in hypoxia status prior therapy. Development of non-invasive MR imaging to assess hypoxia is crucial in determining the effectiveness of TH-302 therapy and to follow response. In further studies, our approach can be used to better plan therapeutic schedules to avoid resistance.

**One Sentence Summary:** Development of non-invasive MR imaging to assess hypoxia is crucial in determining the effectiveness of TH-302 therapy and to follow response.

## Introduction

Sarcomas constitute a large and heterogeneous group of malignant tumors of mesenchymal origin, divided into two categories: soft tissue sarcomas (STS) and sarcomas of bone (*1*). Although STS account for only 1.5% of all malignant tumors in adults, with an estimated 13,130 new cases in the United States in 2020, they represent approximately 7.4% of all tumors in children and young adults (*2, 3*).

The heterogeneity of sarcomas is significant with at least 50 different histologic subtypes, all of which have distinct biologic behavior and response to therapy (*4*). Rhabdomyosarcoma is the most common STS histological type in children. While it is typically initially sensitive to chemotherapy initially, a durable control of the primary tumor requires surgical resection and/or radiation therapy (*5, 6*). Fibrosarcoma is rarer, currently accounting for 3.6% of STS (*7*). Regardless the subtype, however, the standard of care for non-rhabdomyosarcoma STS patients is fairly homogeneous: first-line chemotherapeutic agents, such as doxorubicin (Dox), surgery, and radiation. However, the clinical response to these drugs is limited (*8*).

Sarcoma often presents with significant tumor hypoxia, which has been associated with biochemical resistance to chemo- and radio-therapies and poor prognosis (*9*). Hypoxic tissue also associated with poor vascular perfusion, which can lead to inefficient drug delivery and hence physiological resistance (*10*). Hence, regional hypoxia can subsequently lead to the formation of localized environmental niches where drug-resistant cell populations can survive, evolve, and thrive. Thus, targeting hypoxia in the tumor microenvironment is of great interest to improve clinical outcome (*11*).

For this purpose, hypoxia-activated prodrugs (HAPs) have been designed to penetrate into hypoxic regions and release cytotoxic agents. Multiple HAPs are in development with low toxicity and proven efficacy in pre-clinical and early stage clinical trials (*12*). Evofosfamide (TH-302) is a HAP created by linking a 2-nitroimidazole moiety to the DNA cross-linker bromo-ifosfamide mustard (Br-IPM). As with most HAPs, TH-302 is selectively reduced under hypoxic conditions, which releases the Br-IPM leading to DNA crosslinking (*13*). HAPs have been tested in clinical trials but despite early promise in early phase I-II trials, definitive phase III trials have failed to show a survival benefit (*14*). Specifically, the randomized two-arm phase III clinical trial TH CR-406/SARC021 (NCT01440088) tested TH-302 + Dox *vs*. Dox in patients with locally advanced, unresectable, or metastatic soft-tissue sarcomas. Although the proportion of patients who achieved complete or partial response was significantly higher and progression free survival (PFS) was prolonged for the TH-302 + dox arm, this combination did not improve overall survival (OS) when compared with Dox monotherapy, which was the primary endpoint (*15*). Some limitations have been discussed regarding the lack of OS improvement in this trial (*16, 17*), but perhaps the most important limitation was the lack of patient stratification based on hypoxic status to identify patients who were most likely to benefit from HAP therapy and simultaneously least likely to benefit from Dox monotherapy (*18*).

In this context, we propose that imaging of hypoxia may help with patient stratification and therapy monitoring (*19-22*). To this end, different magnetic resonance imaging (MRI)- and positron emission tomography (PET)-imaging based analyses have been explored to identify tumor hypoxia (*23, 24*) and predict response to HAPs in pre-clinical models (*19, 25*). We have previously reported that multiparametric (mp) MRI can capture subtle difference in the tumor microenvironments, and is able to differentiate viable, necrotic and hypoxic tumor habitats in breast cancer models (*26*).

The main goal of this study was to use multiparametric MRI (mpMRI) and co-registered histology to classify hypoxic habitats, in order to monitor therapy response to TH-302 in preclinical models of sarcoma. Thus, this study had two specific goals: first, to evaluate the response to TH-302 monotherapy or in combination with Dox in sarcoma mouse models; and second to develop a combined Deep-Learning and MRI-based method to identify hypoxic habitats to investigate the temporal evolution of changes in hypoxic habitats noninvasively in sarcomas over the course of treatment.

## Results

### Tumor growth and survival

Two sarcoma models were used in this study: a patient-derived xenograft (PDX) of rhabdomyosarcoma and a murine fibrosarcoma syngeneic model (RIF-1). Therapies with dox, TH-302 or TH-302 + dox were initiated for each mouse when their tumors reached approximately 500 mm^3^ (day 0). Tumor volumes and MRI was performed until tumors individually reached a volume of ≥ 1500 mm^3^, at which point animals were terminated.

In the <u>rhabdomyosarcoma PDX model</u>, monotherapy with TH-302 or the combination of Dox + TH-302 resulted in reduced tumor growth, while dox monotherapy was not effective **(Fig. 1A)**. Compared to untreated control or the Dox monotherapy arm, the OS significantly increased with both the TH-302 monotherapy (p=0.0019 *vs* Control; p=0.0016 *vs* Dox) and the Dox + TH-302 combination (p=0.0046 *vs* Control; p=0.0035 *vs* Dox). The median survival for control and Dox-treated groups were 9 and 7 days respectively, while it increased to 35 and 82 days when mice were treated with TH-302 monotherapy or TH-302+Dox combination, respectively (**Fig. 1B)**. The superiority of combination therapy in increasing OS over the TH-302 monotherapy was consistent with concept that TH-302 controls hypoxic habitats while an anti-proliferative agent, such as Dox, controls the normoxic viable tumor areas (*27*). In the combination group, 4 of 5 tumors regressed during therapy, but eventually regrew (**Fig. 1A**), suggesting that they may have acquired a resistance mechanism during prolonged therapy. All therapies were well tolerated, and mice did not show significant changes in body weight during therapy (p-values > 0.05; **Fig. S1)**.

**Fig. 1.**
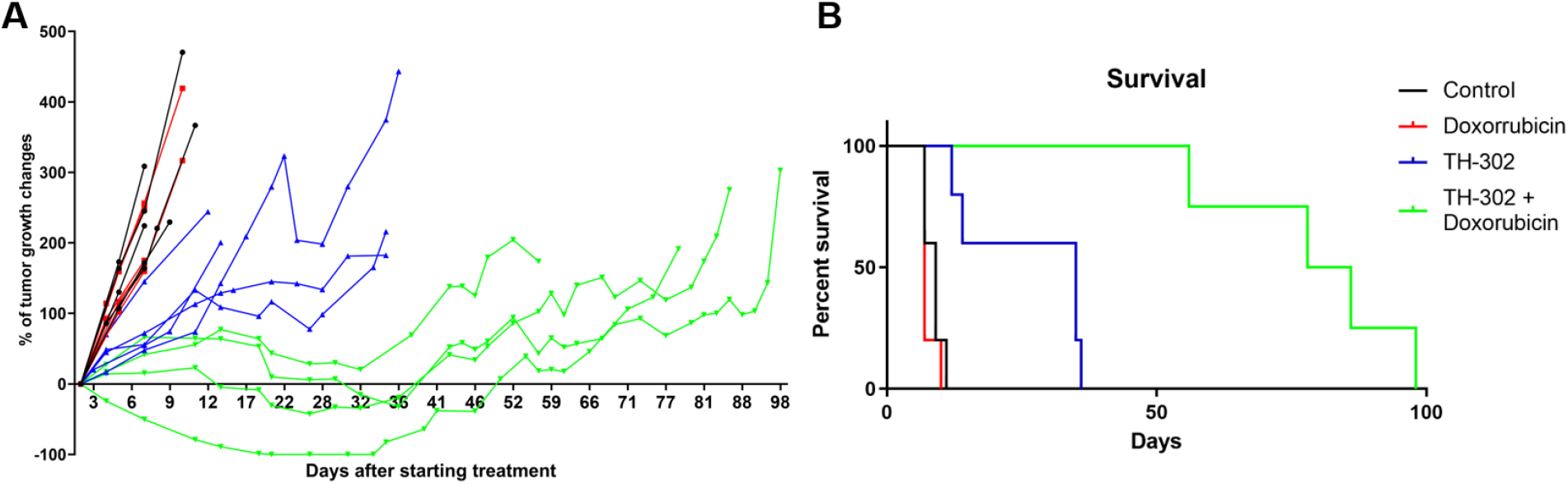
Tumor growth and survival plots for patient-derived xenograft (PDX) rhabdomyosarcoma model. **A**. Tumor growth changes (%) after starting treatment (day 0). **B**. Kaplan-Meier plots shown that monotherapy with TH-302 or with the TH-302 + doxorubicin combination increased the overall survival (p=0.35 Dox *vs* Control; p=0.019 TH-302 *vs* Control; p=0.0016 TH-302 *vs* Dox; p=0.0046 TH-302 + Dox *vs* Control; p=0.0035 TH-302 + Dox *vs* Dox; p=0.0051 TH-302 + Dox *vs* TH-302 + Dox).

In the <u>Radiation-Induced Fibrosarcoma (RIF-1)</u> model, no therapy had a significant effect on the tumor growth rates or survival. There were no differences in the tumor growth between the control mice and mice treated with any therapy **(Fig. 2A)**. In addition, there were no differences in the OS, with median survivals of 5; 5; 7 and 6.5 days for the Control, Dox, TH-302, or Dox + TH-302 groups, respectively (p=0.71; p=0.08; p=0.16; **Fig. 2B)**. Mouse body weight was not affected during any therapy protocol (p-values >0.05; **Fig. S2)**.

**Fig. 2.**
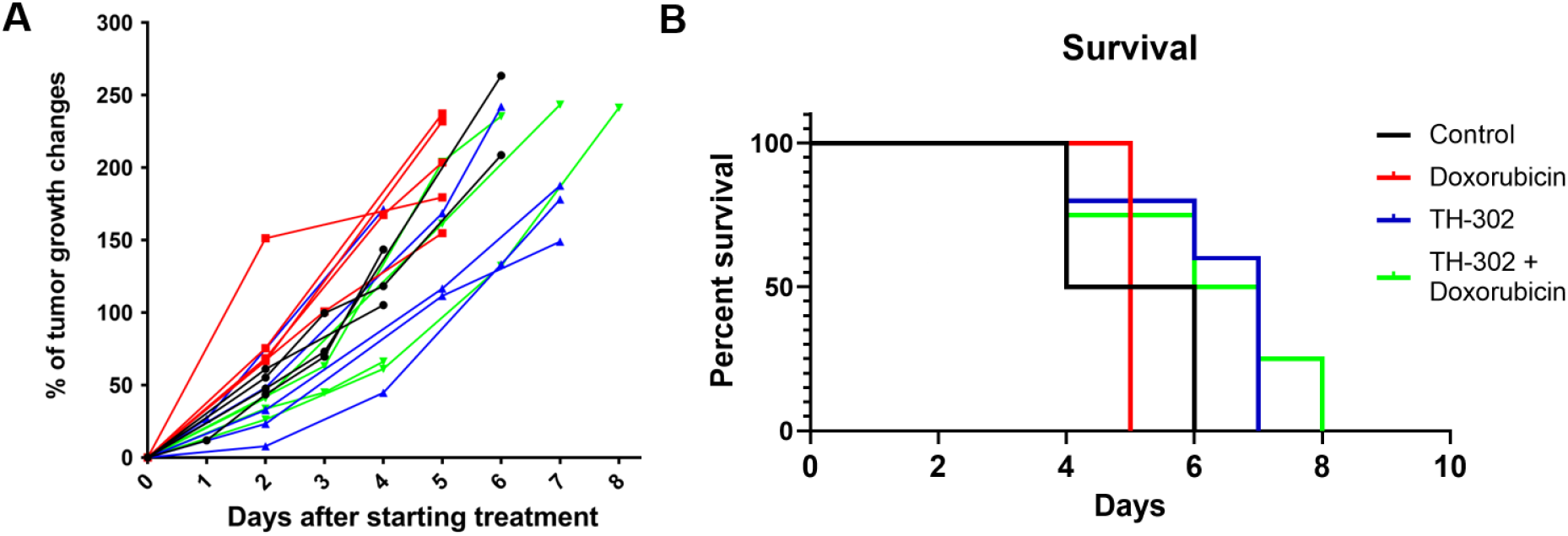
Tumor growth and survival plots for Radiation-induced fibrosarcoma cell line (RIF-1) model. **A**. Tumor growth changes (%) after starting treatment. **B**. Kaplan-Meier plots shown that there was not significant difference in the OS between groups of treatment (p=0.13 Dox *vs* Control; p=0.08 TH-302 *vs* Control; p=0.06 TH-302 *vs* Dox; p=0.16 Dox + TH-302 *vs* Control; p=0.12 Dox + TH-302 *vs* Dox; p=0.73 TH-302 *vs* Dox + TH-302).

### Hypoxia status can determine TH-302 response

As hypoxia is the main determinant for TH-302 activity, we evaluated the hypoxic status of RIF-1 and PDX tumors using pimonidazole (PIMO) staining. PIMO is an exogenous 2-nitroimidazole probe that binds covalently to thiol-containing proteins when the O_2_ tension is below 10 mmHg (< 1.3%) (*28*) and can be visualized in histological sections by immunohistochemistry. For this study, tumors were collected at the time of sacrifice and PIMO-staining was quantified and compared between groups. Surprisingly, the percentage of PIMO-positive area was statistically higher in the last day of therapy in the PDX tumors treated with Dox + TH-302 combination when compared with control (p=0.006) and Dox-treated tumors (p=0.005) (**Fig. 3A)**. The TH-302 monotherapy treated-tumors also appeared to have increased hypoxia, but the results were not significant (p=0.36). For RIF-1 tumors, there was no significant difference in PIMO-positive areas between control and any of the therapy groups **(p>0**.**05; Fig. 3B**).

**Fig. 3.**
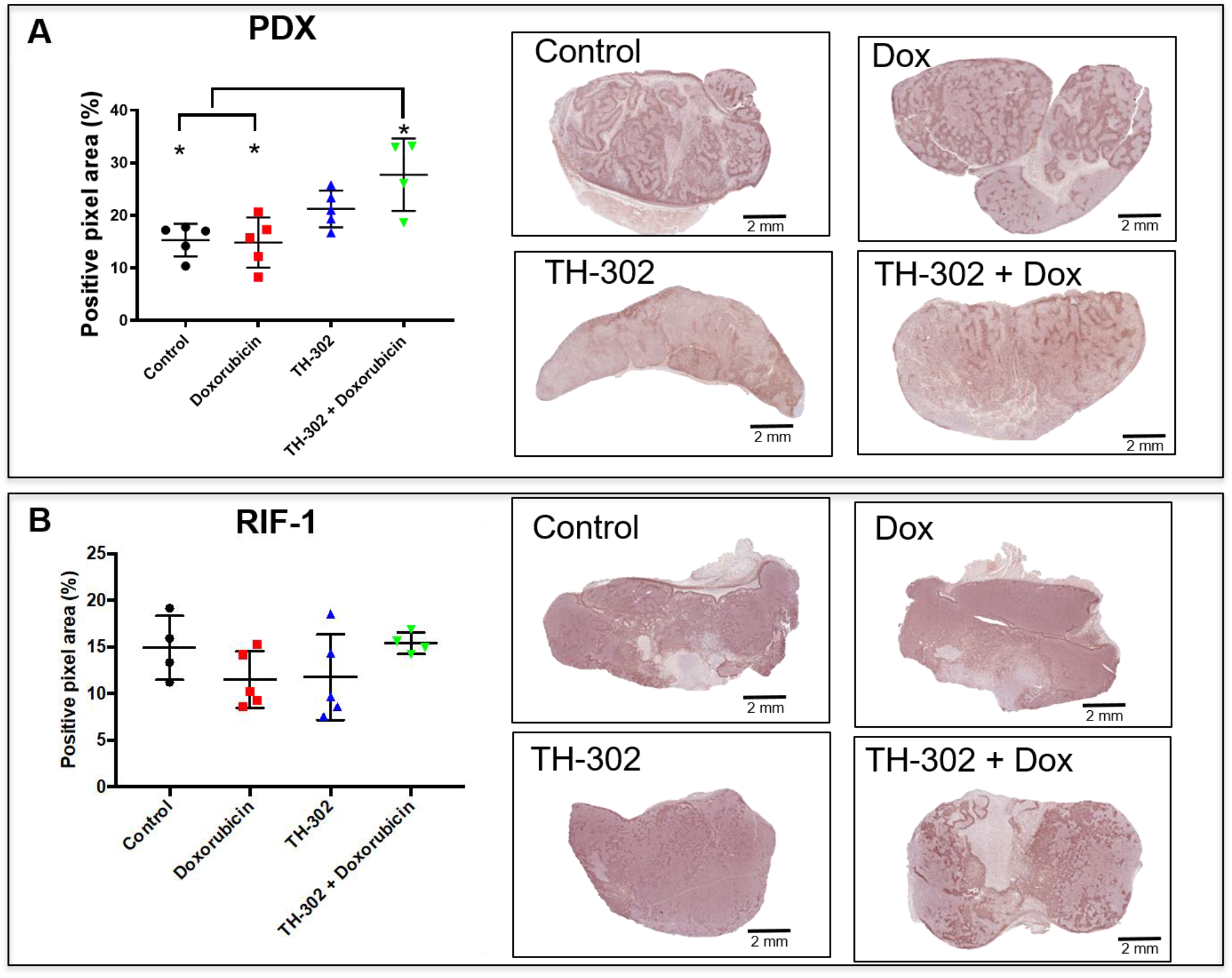
Quantification and representative images of pimonidazole staining in tumors collected in the last day of therapy. **A**. PDX rhabdomyosarcoma model. (p>0.99 Dox *vs* Control; p=0.36 TH-302 *vs* Control; *p=0.006 TH-302 + Dox *vs* Control; p=0.27 TH-302 *vs* Dox; *p=0.005 TH-302 + Dox *vs* Dox; p=0.32 TH-302 *vs* TH-302 + Dox). **B**. RIF-1 model; (p=0.94 Dox *vs* Control; p>0.99 TH-302 *vs* Control; p>0.99 TH-302 + Dox *vs* Control; p>0.99 TH-302 *vs* Dox; p=0.65 TH-302 + Dox *vs* Dox; p=0.78 TH-302 *vs* TH-302 + Dox). ANOVA followed by Bonferroni multiple comparison test. Values presented as mean± SD.

In order to further investigate the different responses to TH-302 in PDX and RIF-1 models, tumors were also stained with a blood vessel marker CD31. RIF-1 tumors were much more vascularized than PDX tumors, showing significantly higher CD-31 staining **(p=0**.**0002; Fig. S3)**, suggesting that a well-perfused and well-oxygenated tumor environment can be contributing to the non-response to TH-302 in the RIF-1 tumors.

### RIF-1 resistance to TH-302 therapy is due to lack of hypoxia

To further explore the mechanisms responsible for TH-302 resistance in RIF-1 model, we evaluated if immune response present in the syngeneic model (RIF-1 cells inoculated into CH3 mice), but impaired in the PDX model (immunodeficient Sho/Scid athymic mice) could be contributing to the TH-302 resistance. To do this, we grew RIF-1 tumors in NSG (NOD scid gamma mouse) immunodeficient mice. As shown in **Fig. S4**, TH-302 therapy was not effective in this immunocompromised RIF-1 model, showing similar results as observed for the immunocompetent RIF-1 model, suggesting that TH-302 resistance is not mediated by an immune response.

Another possible source of resistance could be a biochemical resistance to alkylating agents possibly through enhanced DNA repair processes. To investigate this, we tested if RIF-1 cells *in vitro* were affected by a DNA cross-linking agent Mitomycin-C (MCC). We used MCC because Br-IPM is too hydrophilic to diffuse at significant rates across the plasma membrane and it is less cytotoxic when added to extracellular medium compared to when it is generated intracellularly from the prodrug (*29*). As shown in **Fig. 4**, RIF-1 cells viability was highly sensitive to MCC, indicating that these cells can respond to alkylating agents, such as Br-IPM. H460 cells (human lung cancer cell line) were used as positive control as a known MCC sensitive line.

**Fig. 4.**
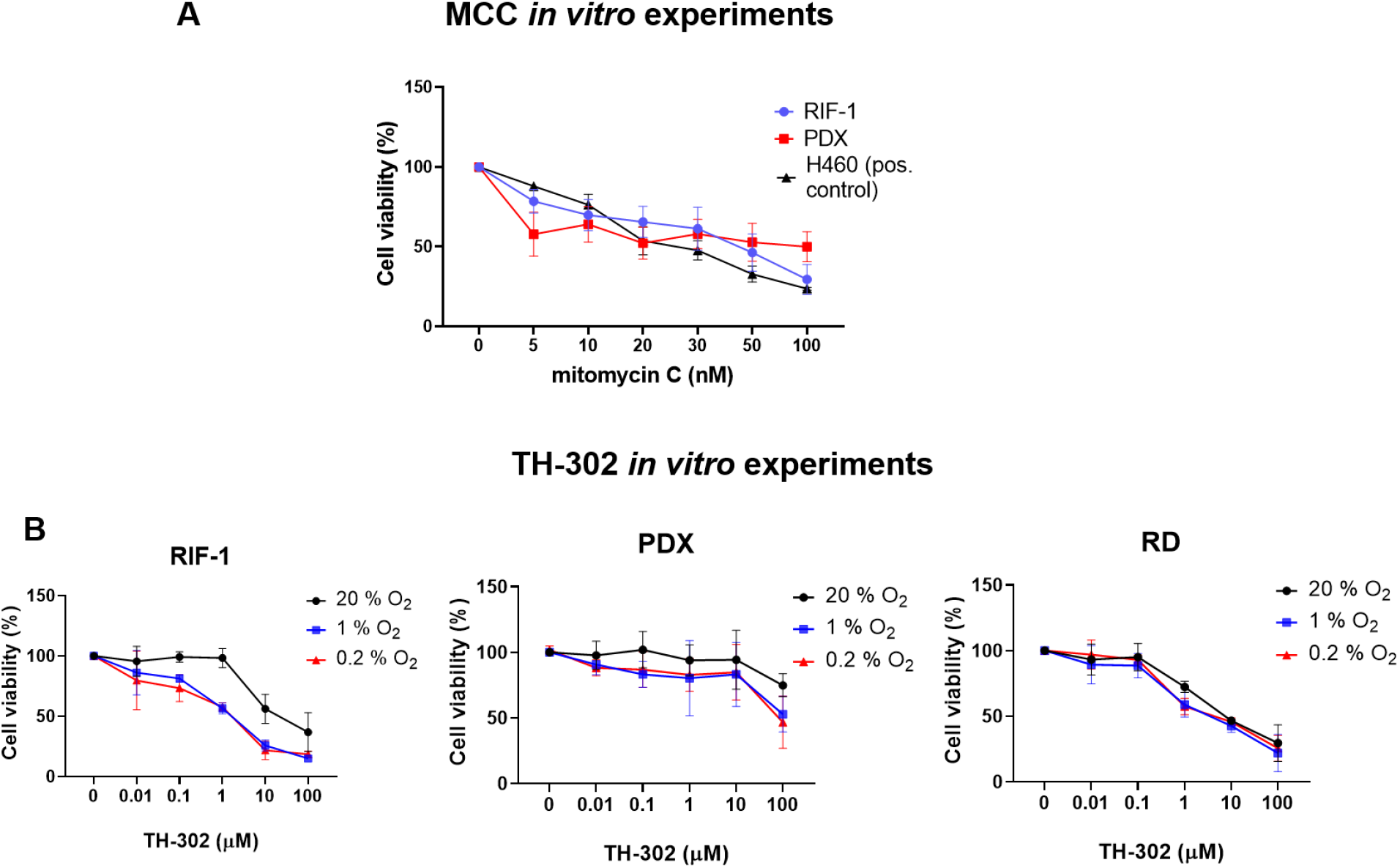
*In vitro* experiments to test cell viability (%). **A**. Dose-dependent mitomycin C (MCC) treatment for 72 hours. Only p-values <0.05 are shown. P-values for RIF-1 cells: *p=0.007 for Control (C) (0 nM) *vs* 50 nM MCC; *p=0.0007 for C *vs* 100 nM MCC. P-values for PDX cells: *p=0.04 for C *vs* 100 nM MCC. P-values for H460 cells: *p<0.001 C *vs* ≥ 20 nM MCC. **B**. Dose-dependent TH-302 treatment under normoxic (20% O2) or hypoxic conditions (1% O2 and 0.2% O2). Only p-values <0.05 are shown. P-values for RIF-1 cells: *p<0.001 for Control (C) (0 µM) *vs* ≥ 10 µM at 20% O2, and ≥ 1 µM at 1% O2 and 0.2% O2. p-values for PDX cells: *p<0.01 for C *vs* 100 µM at 1% O2 and 0.2 % O2. p-values for RD cells: *p<0.005 for C *vs* ≥ 1 µM at 20% O2, 1% O2 and 0.2% O2. ANOVA followed by Dunnett’s multiple comparisons test.

Finally, to check if controlled conditions of hypoxia would improve sensitivity to TH-302 in the RIF-1 cells, we tested TH-302 therapy *in vitro* under hypoxia and normoxia. Of note, RIF-1 cells were strongly sensitive to TH-302 under hypoxic conditions. There was a concentration-dependent response to TH-302 under hypoxia, while only higher concentrations were effective under normoxia in both the RIF-1 and PDX cells. RD cells (human rhabdomyosarcoma cell line) were used as positive control (*30*) and showed sensitivity to TH-302 in doses > 1 µM under normoxia and hypoxia conditions (**Fig. 4)**.

### Noninvasive measurement of hypoxia in MR imaging

The above data indicates that RIF-1 cells can respond to TH-302, but only at hypoxic conditions. It emphasizes the importance of identifying tumor hypoxia at the stage of treatment planning to therefore predict response.

In addition, increase of PIMO staining observed in TH-302 and TH-302 + Dox treated tumors collected on the last day of therapy indicate that there is a substantial amount of hypoxia prior to and following HAP therapy. This raises the question whether there were changes in hypoxia during therapy. PIMO is often used as the “gold standard” to measure hypoxia in tissues, but a major limitation is that it requires collection of tissue for histology, which is not conducive for longitudinal measurements. Although a biopsy can be taken to assess hypoxic status prior to or during therapy, it would not account for intratumor heterogeneity and would be prone to interfering with the study. In this context, if noninvasive imaging of hypoxia can be developed it would allow longitudinal monitoring of therapy response without a biopsy. We had previously developed a multiparametric MRI (mpMRI) method to identify hypoxic habitats in breast cancers using Gaussian Mixture Models (*26*). Herein, we take a similar approach to identify hypoxia in sarcoma using a convolutional neural network (CNN).

This network was trained using co-registered PIMO-stained histology and MRI slices. Histological slices stained with PIMO were co-registered with MRI slices using 3D-printed cradle as described in Methods and DICE similarity coefficients (DSC) between contour-defined areas of the MR and histology images were calculated. For the PDX tumor model, 57 slices were co-registered, and the average of DSC scores was 0.92 ± 0.02 (median = 0.93) while for the RIF-1 tumor model 63 slices were co-registered and average of scores was 0.93 ± 0.02 (median = 0.93) **(Tables S1 and S2**). Thus, only samples with greater than average similarity scores of ≥0.92 were used to develop the CNN models.

Based on mouse-level data with a preferred training/validation/testing ratio of 60/10/30; 43 slices of 18 PDX models were randomly divided into a training (n=25), validation (n=4) and test datasets (n=14), and 49 slices of 15 RIF-1 models were randomly divided into a training (n=26), validation (n=5) and test datasets (n=18). A cutoff of 0.4 was used to binarize the CNN predicted hypoxia probability for both PDX tumors and RIF-1 tumors to obtain the predicted hypoxic habitats. The mpMRI maps, co-registered PIMO stained histology slice and the predicted portion of hypoxia of PDX and RIF-1 tumor models from training, validation and test datasets are shown in **Figs. 5 and 6**, respectively.

**Fig. 5.**
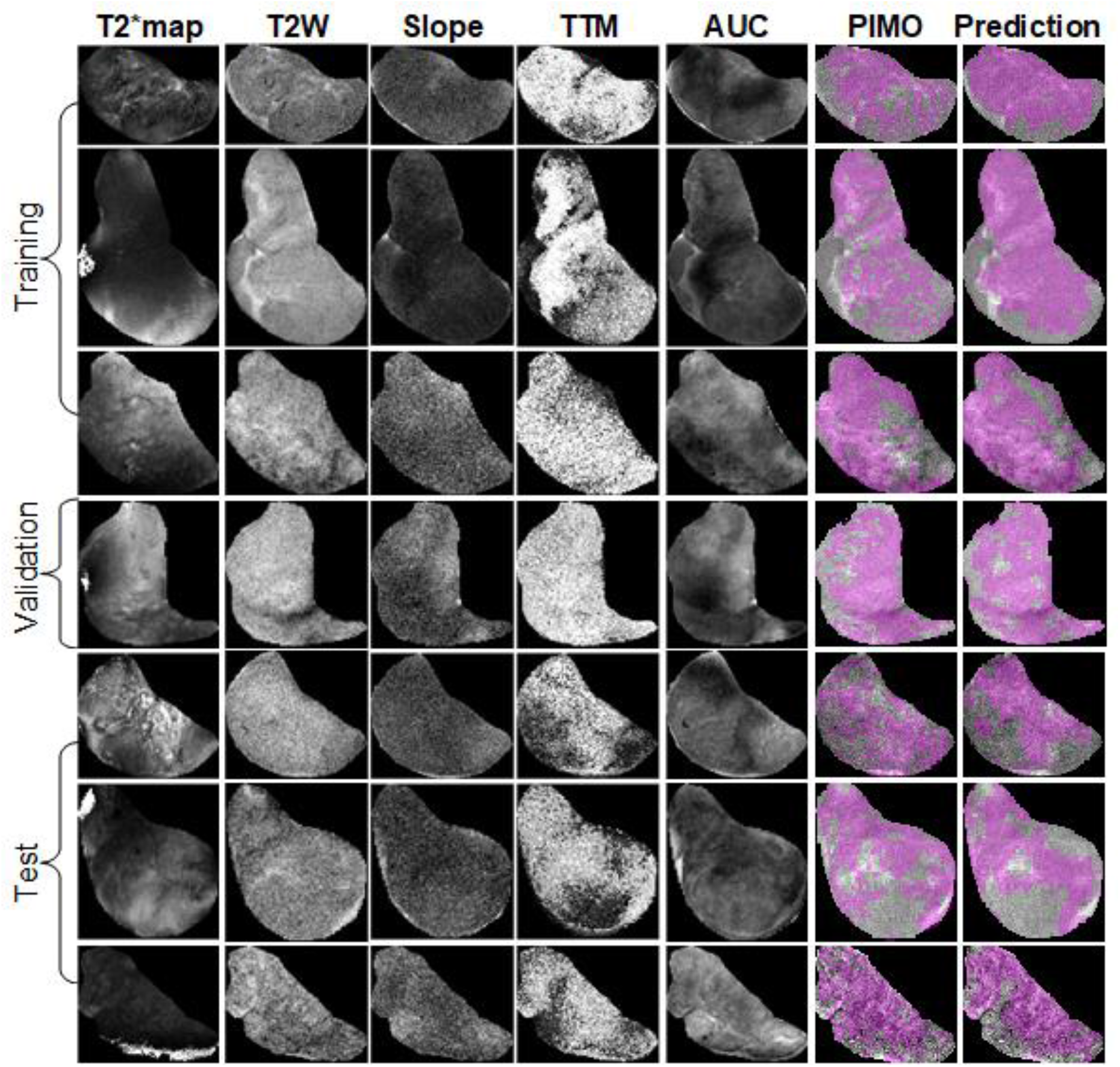
Representative samples of training, validation and test datasets for PDX tumors, showing the mpMRI maps: T2star (T2*) map, T2-weighted image (T2W) and slope, time to maximum (TTM), and area under the time-series curve (AUC) from Dynamic contrast enhanced (DCE) MRI, co-registered pimonidazole stained histology slice (PIMO), and predicted portion of hypoxia (Prediction).

**Fig. 6.**
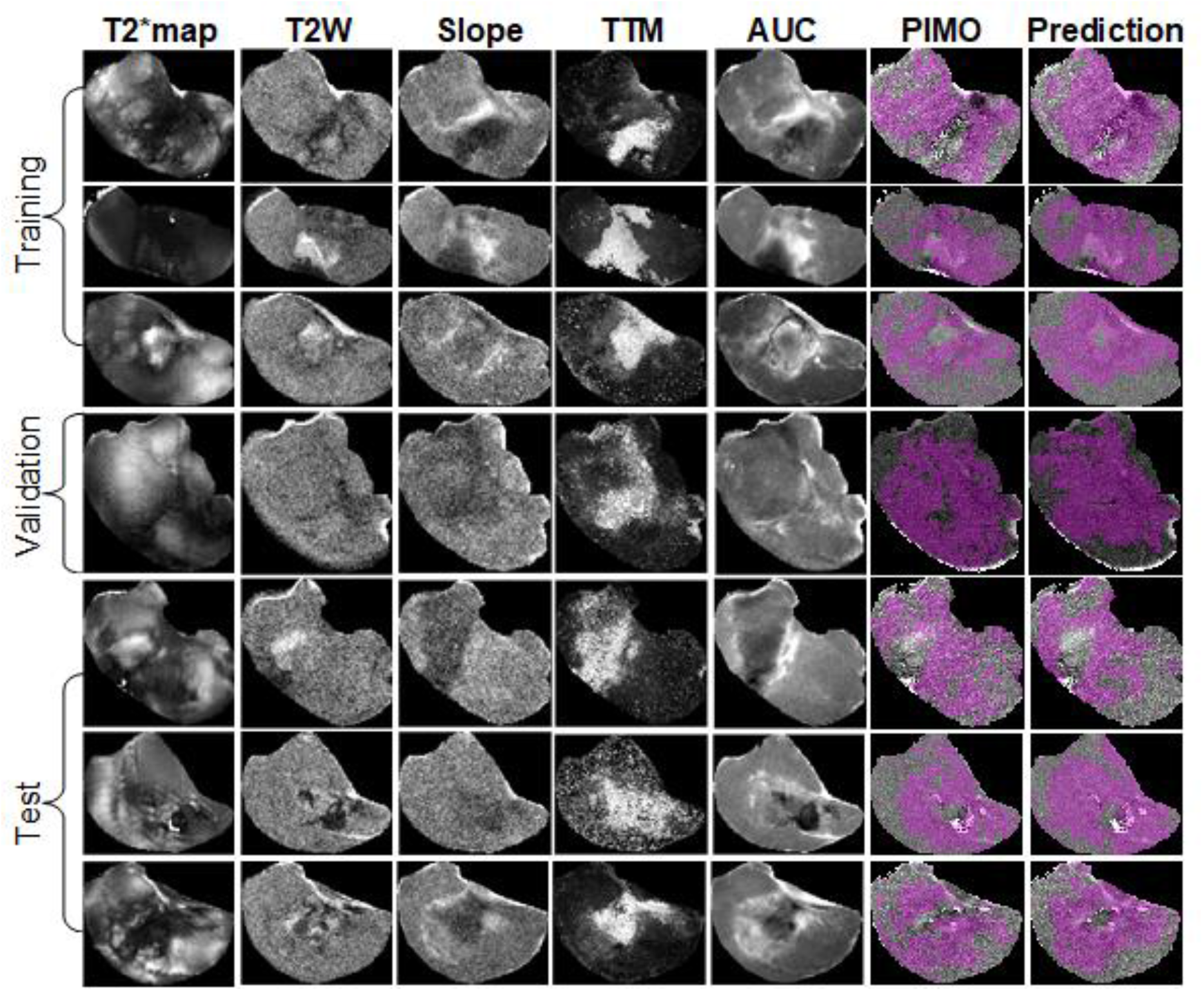
Representative samples of training, validation and test datasets for RIF-1 tumors, showing the mpMRI maps (T2star (T2*) map, T2-weighted image (T2W) and slope, time to maximum (TTM), and area under the time-series curve (AUC) from Dynamic contrast enhanced (DCE) MRI, co-registered pimonidazole stained histology slice (PIMO), and predicted portion of hypoxia (Prediction).

For PDX tumors, a strong correlation of 0.82 (p<0.001), 0.82 (p=0.18), and 0.81 (p<0.001) was found between true positive portion and predicted positive portion in the training, validation and test cohorts, respectively. For the RIF-1 tumors, the correlation was also as strong at 0.85 (p<0.001), 0.93 (p=0.023) and 0.76 (p<0.001) in the training, validation and test cohorts, respectively. Detailed plots are provided in **Fig. S5. Tables S3 and S4** show detailed quantitative metrics for each slice.

### Hypoxia status prior to therapy can determinate TH-302 response

Using these models, we were able to non-invasively identify and quantify hypoxia longitudinally and at pre-therapy MR imaging. Comparison of hypoxic status at pre-therapy between tumor models confirmed that “predicted portion of hypoxia” was significantly lower in RIF-1 than PDX tumors, which is consistent with the response to TH-302 in the PDX model, and resistant in the RIF-1 model (**Fig. 7**).

**Fig. 7.**
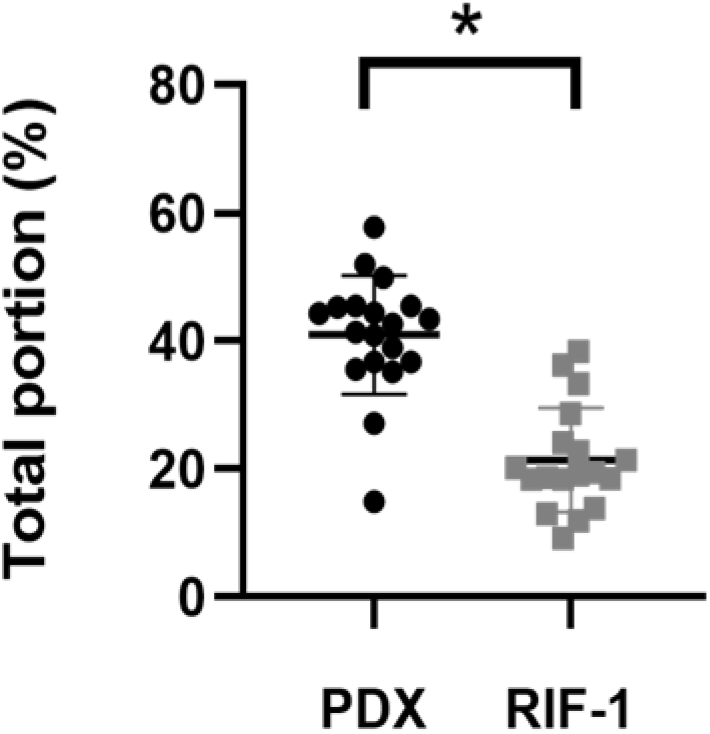
Comparison between predicted positive portion of hypoxia in pre-therapy MR imaging for PDX and RIF-1 tumor models. *p<0.0001, Student t-test.

This data corroborated with the *in vitro* findings which shows that RIF-1 are sensitive to TH-302 only under hypoxic conditions. Once again, it reinforces the need of measuring hypoxic status prior therapy to determinate response to TH-302.

Interestingly, survival of PDX mice treated with TH-302 therapy monotherapy increased to 35 days, however there was one mouse that did not respond to TH-302, reaching the limit tumor volume 12 days after starting therapy. In fact, this mouse showed the lowest levels of predicted hypoxia in the pre-therapy MRI for this group, which is consistent with the non-response to TH-302.

To test if levels of hypoxia prior to therapy could predict response to TH-302, we analyzed all mice treated with TH-302 and TH-302 + Dox regardless of PDX or RIF-1 tumor models, consistent with the SARC21 (NCT01440088) clinical trial wherein STS patients were treated regardless of histotype. Here, we asked whether the “predicted portion of hypoxia” at pre-therapy imaging could predict the response to TH-302 (survival days ≥ 14 days) with an AUROC of 0.89 (95%CI: 0.72, 1.00, p=0.005), and also achieved C-index of 0.73 (95%CI: 0.57, 0.88, p=0.004) in predicting the survival. The optimal cutoff of 25.60% was obtained based on the maximum Youden index in the ROC curve to identify the mice which are more likely to respond to TH-302. Using this cutoff, mice were stratified into high- and low-hypoxia portion groups. The mice within the high-hypoxia portion group had longer survival days with median value of 35 (interquartile range (IQR): 14-75) days versus 7 (IQR: 6-7) days of the low-hypoxia portion group. Using Cox proportional hazards regression analysis, the binarized predicted portion of hypoxia with this cutoff was identified as significant prognostic factor with the hazard ratio (HR) of 0.27 (95%CI: 0.090, 0.83, p=0.022) in survival prediction.

### Temporal evolution of hypoxic habitats

**Fig. 8A-F** showed longitudinal measurements of “predicted portion of hypoxia” for the PDX tumor model. It is possible to observe an increase of hypoxia after day 7 for most of the tumors regardless the therapy **(Figs 8A-D**). However, for tumors treated with TH-302 or with TH-302 + Dox, hypoxic portion was decreased or controlled during the course of the therapies (**Figs. 8C-D**). More specifically for the TH-302 group, one mouse showed a decrease of 9.5% at day 7, while it was decreased or controlled in other 3 mice after day 14. For the mouse that did not respond to TH-302, there was an increase of 31% of hypoxia from pre-therapy to day 12 (black diamond symbol in **Fig. 8C**). All mice treated with TH-302 + Dox showed a decrease in hypoxia at different time points during therapy. In the first measurement after starting therapy, hypoxic portion decreased in one mouse but increased in the other 3 mice. It was reduced at day 14 for one mouse, and after day 50 for the other 2 mice, however, eventually an increased in hypoxia was observed in this group, as the tumors grew, and the therapies’ effectiveness were reduced (**Fig 8D**).

**Fig. 8.**
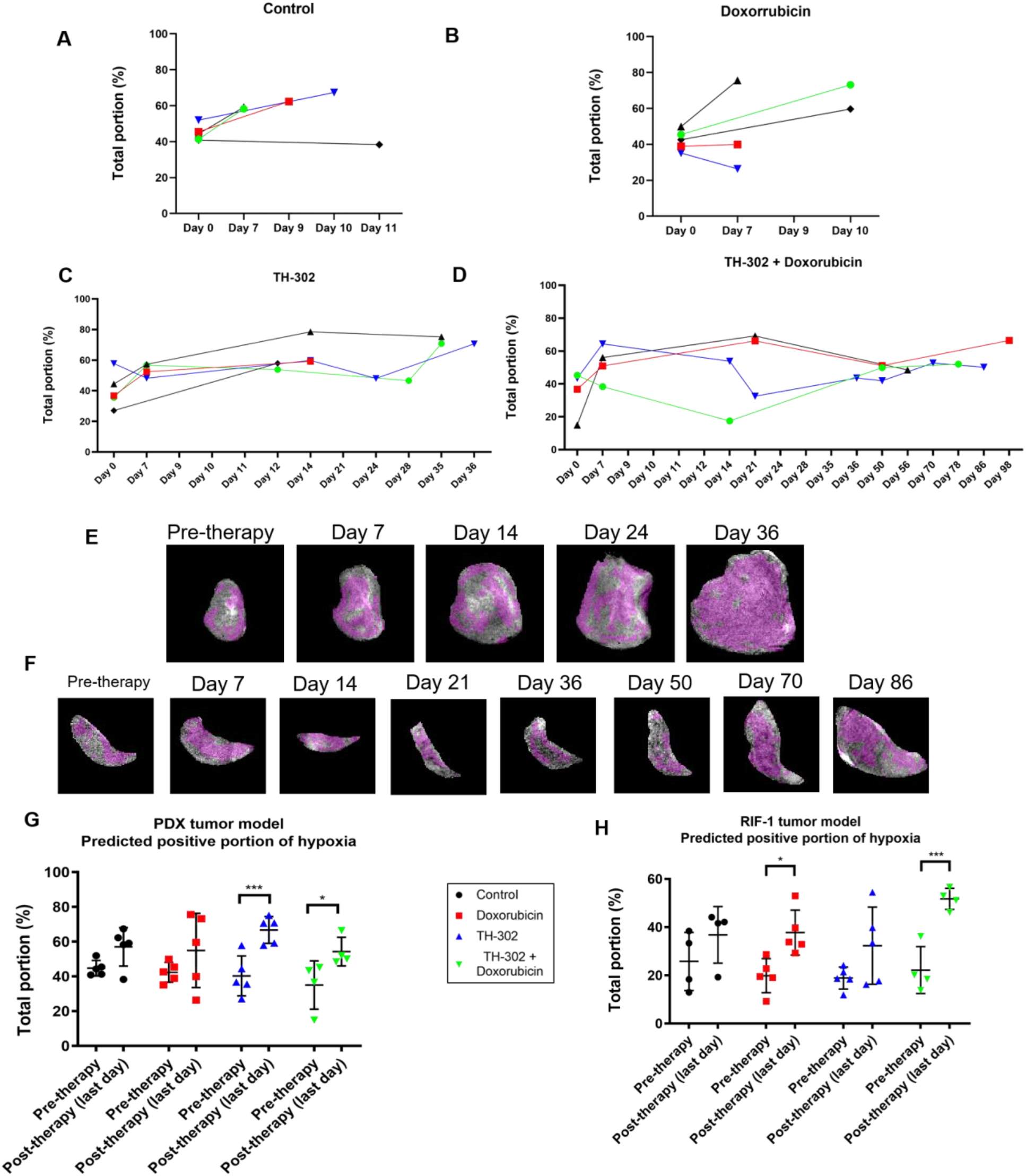
Longitudinal measurements of predicted portion of hypoxia in MRI imaging for the PDX tumor model. **A**. Control group; **B**. Dox treated group; **C**. TH-302 treated group; **D**. TH-302 + Dox treated group; **E**. Representative images of changes in predicted portion of hypoxia (in magenta) during therapy for a TH-302-treated tumor; **F**. Representative images of changes in predicted portion of hypoxia (in magenta) during therapy for a TH-302 + Dox-treated tumor; **G**. Predicted portion of hypoxia in pre-therapy and last day of therapy MR imaging for the PDX tumor model (Pre-therapy *vs* last day: p=0.15 Control; p=0.14 Dox; ***p=0.0009 TH-302; *p=0.02 TH-302 + Dox). **H**. Predicted positive portion of hypoxia in pre-therapy and last day of therapy MR imaging for the RIF-1 tumor model. (Pre-therapy *vs* last day: p=0.35 Control; *p=0.01 Dox; p=0.10 TH-302; ***p=0.0009 TH-302 + Dox). ANOVA followed by Bonferroni test.

When comparing values of “predicted portion of hypoxia” from pre-therapy with the final measurement taken before sacrifice, it was observed that there appeared to be more hypoxia in the last day of therapy for all groups of PDX tumors. Unexpectedly, this increase in hypoxia was statistically significant in the tumors treated with TH-302 monotherapy (p=0.0009) and with the TH-302 + Dox combination (p=0.02) (**Fig. 8G)**. For RIF-1 tumors, the predicted hypoxic portions were also higher on the last day of therapy, and it was statistically significant for the dox-treated tumors (p=0.01) and tumors treated with the TH-302 + dox combination (p=0.0009) (**Fig. 8H**). These results are consistent with PIMO staining observed at the histology collected before sacrifice, which showed higher levels in TH-302 and TH-302 + Dox treated groups than control and Dox-treated groups (cf. **Fig. 3**). As discussed below, this increase in hypoxic volume fraction may be due to the tumors’ increased volume, further emphasizing the need to measure hypoxic fractions longitudinally.

We also noted the percentage of cells stained with the apoptosis marker Cleaved Caspase 3 (CC3) in the TH-302 treated tumors was significantly lower than control and dox-treated tumors in the last day of therapy (p<0.05; **Fig. S6A**), while phospho γ-H2AX, a marker for DNA-damage did not show significant changes (p>0.05; **Fig. S6B**), which corroborated the hypothesis that, at the time of sacrifice, tumors were no longer responding to therapy.

## Discussion

In this study, we used the HAP TH-302 to target hypoxia in sarcoma mouse models, and we developed a deeply learned MRI-based method to predict hypoxia prior to and longitudinally during therapy. We showed that TH-302 monotherapy or in combination with Dox was able to delay tumor growth and increase survival in a PDX model of rhabdomyosarcoma, while a syngeneic RIF-1 fibrosarcoma model was resistant to TH-302.

Hypoxic status has been associated with TH-302 response in several pre-clinical models (*19, 27, 31*). However, resistance was observed in hypoxic tumors (*32*), as TH-302 efficacy is also dependent on other conditions, such as prodrug-activating reductases, intrinsic sensitivity to the drug warhead and DNA repair status (*33*). Here, we explored different mechanisms that could be contributing to TH-302 resistance in RIF-1 model and showed that hypoxia status may be the causal effect. Pre-therapy MR imaging showed that RIF-1 tumors are less hypoxic than PDX tumors. In fact, RIF-1 tumors are known to present a small fraction of radiobiologically hypoxic cells, ranging from 0.9% to 1.7% when tumor volume increased from 200 mm^3^ to 2000 mm^3^ (*34, 35*).

This emphasizes the importance of knowing the hypoxia status in order to individualize and hence optimize therapies using HAPs (*14*). Multiple approaches exist to detect hypoxia (*36*), but priority must be given to imaging approaches, which can represent spatially heterogeneous distribution of hypoxia, and are noninvasive and reproducible, allowing not only pre-therapy measurement, but also longitudinal assessment of hypoxia to follow therapy response. This is especially important to optimize combination therapy regimens considering the tumor evolutionary dynamics.

Here, we showed that combination of TH-302 + dox was much more effective than TH-302 monotherapy in the PDX model; however, both therapeutic regimes lead to resistance with prolonged treatment. Higher levels of hypoxia were observed in both these groups at the end of therapy compared to the control and dox-treated groups.

A rationale for using a combination of TH-302 and dox is to target two different populations within the heterogeneous tumor microenvironment, which would lead to complete tumor eradication or long-term control compared to monotherapies that affect only the normoxic or hypoxic adapted populations. However, optimal therapy efficiency is highly dependent on identifying right timing and administration sequence of combination therapies (*11*). In our study, dox was given once a week and TH-302 was given daily, 5 days a week.

Thus, our data suggest that with the continued use of TH-302 and dox, a state of dynamic equilibrium between hypoxic and viable normoxic tumor cell populations was lost, with TH-302 not being able to continue controlling the population in the hypoxic habitat. Therapy-sensitive and resistant cell types constantly compete into the tumor microenvironment; however, this equilibrium can change with prolonged treatment, leading to emergence of a resistant population. The residual cell population that persisted after the several rounds of therapy is likely to have greater intrinsic or environmental resistance, and will continue to survive with the continued use of the same therapeutic regimen (*37, 38*). A different study reached a similar conclusion with an EGFR targeted agent with a mathematical model showing that longer time under TH-302 therapy without the targeted inhibitor erlotinib allowing the EGFR sensitive cell population to expand drastically due to TH-302 resistance. Optimal schedule of an agent affecting both the hypoxic and normoxic populations may allow for the longest duration of control of the two populations (*11*).

In this case, following tumor hypoxia longitudinally in the clinic using imaging approaches as developed in this study, would allow an optimal time planning for switching drugs, avoiding unnecessary doses or drugs and could predict future response or resistance to therapy. In addition, it could guide adaptive therapy, which adjusts the time course of therapy to turn it on and off accordingly, to maintain the sensitive cells populations that will compete and continue to suppress the resistant population (*37*).

Our imaging approach has the potential to also provide information on other tumor habitats, as viable and necrotic regions and this was not addressed in this study. Soft tissue sarcomas are highly heterogenous, even within histological subtypes, and a consistent and accurate identification of these viable and necrotic habitats in histology across different samples is challenging, and specific markers are necessary in addition to morphological characteristics.

Taken together, our results show that different responses to TH-302 in PDX rhabdomyosarcoma and RIF-1 fibrosarcoma is associated with status of tumor hypoxia. Non-invasive MR imaging to identify hypoxia prior to therapy can presage the responsiveness to TH-302 and longitudinally monitor its antitumor effect. Notably, PDX xenografts developed physiological resistance during TH-302 monotherapy or in combination with dox. Specifically, hypoxia imaging developed here can be applied in further studies, where cycle treatments between TH-302 and Dox can be optimized depending on the extent of hypoxic habitats to avoid or delay the emergence of resistance.

## Materials and Methods

### 1. Sarcoma mouse models

Animal experiments were approved by the Institutional Animal Care and Use Committee (IACUC) (Protocol #4778). Two sarcoma models were used in this study: a patient-derived xenograft (PDX) of rhabdomyosarcoma and a murine fibrosarcoma syngeneic model.

To develop the PDX model, cryopreserved cells from a rhabdomyosarcoma patient-derived tumor (reference number: SJRHB010468_X1) were obtained from The Childhood Solid Tumor Network (CSTN) at St. Jude Hospital (*39*). Tumor was established through subcutaneous implantation into the flank of immunodeficient SCID Hairless Outbread (SHO) mice (female, 6–8 weeks of age). Tumors were measured with digital caliper and were passaged into new mice when them reached >1000 mm^3^. Mice were anesthetized with 2% isoflurane delivered in 1.5 L/min oxygen ventilation and tumors were collected, placed in RPMI 1640 culture media (Gibco, Waltham, MA) and dissected to 3 × 3 mm^3^ pieces. Tumors explants were then implanted into new mice with 50% RPMI1640 / 50% Matrigel.

The fibrosarcoma model was developed by inoculating the radiation-induced fibrosarcoma cell line (RIF-1) (*40*) into immunocompetent C3H mice (female, 6–8 weeks of age). RIF-1 cells were kindly provided by Dr. Zaver M. Bhujwalla, Department of Radiology, Johns Hopkins School of Medicine. RIF-1 cells were maintained in Waymouth’s media (Gibco, Waltham, MA) supplemented with 10% fetal bovine serum (FBS), 1 mM of HEPES and 1% of penicillin/streptomycin (Sigma, St. Louis, MO) at 37°C and 5% CO_2_. RIF-1 cells were confirmed to be of mouse origin and no mammalian interspecies contamination was detected for the sample using short tandem repeat (STR) DNA profiling.

Cells were tested free of mycoplasma (MycoAlert Mycoplasma Detection kit; Lonza, Basel, Switzerland). For tumor inoculation, RIF-1 cells were suspended in Hanks’ Balanced Salt Solution (HBSS) media (Gibco, Waltham, MA) and 1× 10^6^ cells were subcutaneously inoculated in the right flank of mice.

All mice were obtained from Charles River Laboratory and housed in a facility under pathogen-free conditions in accordance with IACUC standards of care at the H. Lee Moffitt Cancer Center.

### 2. Groups of treatment

Tumor volumes were measured through volume delineation of T2-weighted MRI images and when tumors reach approximately 500 mm^3^ (day 0), mice were divided into the following treatment groups: 1) Control; 2) monotherapy with Dox, dose of 4 mg/kg by intravenous (IV) injection once a week; 3) monotherapy with TH-302 (obtained from Threshold Pharmaceuticals, Redwood City, CA, USA), dose of 50 mg/kg by intraperitoneal (IP) injection, five times per week; and 4) combination of Dox + TH-302.

The percentage of tumor growth changes and OS were calculated from the first day of treatment (day 0) until the last of experiment, when individual tumors reached approximately 1500 mm^3^.

For the PDX model, a total of 22 SCID/SHO mice were studied with 3 mice dying during the experiment in the MRI scanner, and were excluded from the study. Treatment groups were composed of 5; 5; 5 and 4 mice in Control; Dox; TH-302; and Dox + TH-302 groups, respectively. For the RIF-1 model, 20 CH3 mice were inoculated with tumor cells, while one mouse did not develop tumor and one died during MRI scanning and was excluded from the study. Treatment groups were composed of 4; 5; 5; and 4 mice in Control; Dox; TH-302; and Dox + TH-302 groups, respectively. In addition, 5 immunodeficient NSG mice were used in an additional experiment, where RIF-1 cells were inoculated into those mice, 2 were used as control and 3 treated with TH-302 monotherapy.

### 3. Cell viability in vitro

Cell viability was assessed by crystal violet to test the *in vitro* response to TH-302 and to a DNA cross-linking agent mitomycin C (MCC). Experiments were performed with RIF-1 cells and Rhabdomyosarcoma PDX-derived dissociated cells, as well as with cell lines of human rhabdomyosarcoma (RD) and human lung cancer (H460), which were used as positive controls.

To obtain the cells from PDX tumors for the *in vitro* experiments, tumors were harvested and enzymatically disassociated using the Animal free Collagenase/Dispase Blend II reagent (Millipore, Burlington, MA). Cells were maintained in culture with RPMI-1640 media (Gibco, Waltham, MA) supplemented with 10% FBS and 1% of P/S at 37°C and 5% CO_2_.

For the cell viability assay, cells were seeded into 96-well plates and grown overnight prior to initiating treatment. On the day of the test, cells were exposed to increasing concentrations of TH-302 and the plates were incubated overnight under normoxic (20% O_2_) or hypoxic conditions (0.2% O_2_ and 1% O_2_). After overnight exposure, plates were removed from the hypoxia chamber and further incubated for 72 hours in standard incubator (20% O_2_). For MCC experiments, cells were treated with different concentration of MCC for 72 hours under normoxia (20% O_2_).

### 4. MRI

#### 4.1. Tumor volumes measurements

Tumor volumes were periodically measured by acquiring multi-slice axial T2-weighted MR images covering the entire tumor (TurboRARE sequence; Repetition time (TR) = 4825 milliseconds (ms), Echo time (TE) = 73 msfield of view (FOV) = 35 × 35 mm^2^, matrix = 256 × 256, slice thickness of 1 mm). Tumor volumes were obtained from manually drawn regions of interest (ROIs) in MATLAB (Mathworks, MA, USA) using AEDES (aedes.uef.fi).

During MRI scanning, mice were maintained anesthetized with 2% isoflurane delivered in 1.5 L/min oxygen ventilation, and body temperature and respiratory function were continuously monitored (SA Instrument Inc System 1025, Stony Brook, NY).

#### 4.2. Multiparametric MRI (mpMRI)

mpMRI images were acquired pre and post therapy and parameter maps were calculated for T2, T2* and Dynamic contrast enhanced (DCE) MRI. Imaging was acquired for each mouse at day 0 (pre-therapy) and longitudinally until the last day of therapy, when each tumor individually reached a volume of ≥ 1500 mm^3^.

Imaging acquisition and corresponding MR parametric maps were obtained as previously described (*26*). T2 and T2* maps were generated with the multi slice multiecho (MSME) and multi gradient echo (MGE) sequences, respectively. T_1_-weighted DCE-MRI images were acquired pre and post intravenous injection (IV) of 0.2 mmol/kg gadobutrol (Gadavist; Bayer). All sequences were obtained with FOV of 35 × 35 mm^2^, matrix size of 256 × 256, 11 central slices with slice thickness of 1 mm. Images were performed using a 35 mm Litzcage coil (Doty Scientific, Inc) on a 7T horizontal magnet (Agilent ASR 310) and (Bruker Biospin, Inc. BioSpec AV3HD). T_2_ and T_2_* maps were computed in ParaVision (Bruker Biospin, Inc). Semi-quantitative parametric maps calculated from DCE-MRI data included area under the time-series curve (AUC), slope and time to maximum (TTM).

mpMRI maps were co-registered with the corresponding histology and these data were used as input to build DL models to identify hypoxic habitats.

### 5. Histology

#### 5.1. 3D-printed tumor mold

To ensure the co-registration of histology with MRI, tumors were collected after the last mpMR imaging session, according with the 3D-printed tumor mold workflow developed previously (*26*). Briefly, multi-slice axial T2-weighted images were acquired (slice thickness of 1 mm, FOV of 35 × 35 mm^2^ and image size of 256 × 256), and a ROI was drawn encompassing the entire tumor to create a 3D-printed tumor mold. Tumor-specific molds were designed in SolidWorks (Dassault Systems, SolidWorks Corporation, Waltham, MA), containing slots every 2 mm, which were used to guide the slicing of the tumor, aligned with the 1mm MRI slices. Thus, each tumor was sliced in serial 2 mm tissues, placed in individual cassettes and embedded in paraffin to perform immunohistochemistry (IHC) staining. Histology sections were further aligned with the corresponding MRI slices.

#### 5.2. Histological staining

Paraffin tissue-blocks were serially sectioned with slices thickness of 4 µm on a microtome (Leica Byosystem, Buffalo Grove, IL) and allowed to dry at room temperature (RT) and subsequently heated to 60°C for 1 h.

PIMO was used as a hypoxia marker. Mice received an intraperitoneal (IP) injection of PIMO hydrochloride (60 mg/kg) one hour prior to collection of tumors, and PIMO staining was detected by IHC using an anti-PIMO antibody (PAB2627AP, HPI, Burlington, MA). In addition, IHC was performed with the following primary antibodies: Cluster of differentiation 31 (CD31) (#ab28364, Abcam, Cambridge, MA), which was used as a marker of endothelial cells of blood vessels; Cleaved Caspase-3 (CC3) (#9661, Cell Signaling, Danvers, MA) used as an apoptosis marker; and phospho γ-H2AX (#NB100-2280, Novus Biologicals, Littleton, CO) used as a marker for DNA-damage.

IHC protocol consisted in deparaffinization of slides in xylene and hydration through subsequently incubation in aqueous solutions of decreasing ethanol concentration. Endogenous peroxidase activity was blocked with 0.6% H_2_O_2_ in methanol for 30 minutes and antigen retrieval with Citrate Buffer (pH 6.0) in the pressure cooker for 20 minutes. Sections were incubated with 10% goat serum at 4°C overnight for blocking, followed by incubation with primary antibody in humid chamber for 1 hour at RT, and with a biotinylated secondary antibody (Vectastain Elite kit) for 60 minutes at RT. Section were then incubated in Avidin Biotin Complex (ABC) (Vectastain Elite ABC Kit; Vector Labs) for 45 minutes and chromogen substrate visualization was performed using the NovaRed VectaStain Peroxidase kit (Vector Labs SK-4800).

Sections were counterstained with hematoxylin, dehydrated in ethanol followed by xylene, and finally mounted using permount medium (Thermo Scientific, Waltham, MA). Negative controls were obtained by omitting the primary antibody, and a tissue known to express the protein of interest was used as positive controls in every assay.

#### 5.3. Histological analyses

##### 5.3.1. Detection and quantification of positive pixels

Histology slides were scanned (Aperio AT2, Leica Biosystems, Buffalo Grove, IL), saved as .SVS files and imported into Visiopharm software (Visiopharm A/S, Horsholm, Denmark).

First, an intensity threshold algorithm was used to separate the tissue from background and the number of pixels in the tissue area was calculated (total area). Next multiple intensity-based threshold algorithms (pixel intensity from 0 to 255) were created to identify positive stained-pixels for each antibody. A global threshold was used across all images for each antibody (CD31, CC3 and phospho γ-H2AX). Area of positive pixels (%) (stained pixels) was calculated over the total area. The results were verified by the study pathologist.

##### 5.3.2. Binary mask of pimonidazole positive areas

For PIMO staining, Otsu’s method was used to select individual thresholds to mask PIMO-positive pixels in each image. It is known that pimonidazole stains irreversibly when tissue become hypoxic and is a permanent record of when a region has achieved extreme hypoxia. Individual thresholds were chosen to increase accuracy in detecting PIMO positive areas as it was used as true positive portion of hypoxia for the CNN models.

MATLAB was used to automatically select thresholds based on Otsu method for each histological slice. Threshold levels were calculated in grayscale images, on a scale from 0 (no staining, white) to 255 (maximum staining, black). For histological slices of the PDX model, thresholds ranged from 77 to 105 (mean for all slices = 92.06 ± 5.85) and for the RIF-1 slices, thresholds ranged from 74 to 99 (mean for all slices = 86.17 ± 5.58) **(Figs. S7A-B**).

Then, Visiopharm software was used to mask the histological images using the individual Otsu threshold-based algorithms and area of PIMO-positive pixels were calculated. To compare “PIMO-positive area in histology” between groups of therapy, values of all histological slices for each tumor were average to have one value per tumor. Thus, each tumor was represented by a total PIMO-positive area (%).

To ensure that individual thresholding method would not affect the comparison of PIMO-positive areas between groups of therapy, we calculated a global threshold across all PIMO-stained slices for each tumor type. Using these global thresholds (92 for PDX and 86 for RIF-1), we recalculated the PIMO-positive area for each tumor **(Fig. S7C**). Notably, this alternative thresholding method did not significantly affect the values of total PIMO-positive area in each tumor, nor the comparison between groups (**Fig. S8**).

### 6. MRI and Histology co-registration algorithm

Full resolution masks of PIMO-positive pixels were exported from Visiopharm as .mld files and those .mld files were converted into .mat files. Information about positive pixel areas in .mld files is stored as a collection of polygons which are defined by their consecutive vertices. Order of the polygons defines which one encompasses a positive region and which one is a boundary of a negative one (e.g. hole). In order to transform that information into an image of given resolution and store it as a .mat file in MATLAB, each polygon was drawn into an image matrix using Brasenham’s line algorithm and then filled accordingly using queue-scanline algorithm going from top of the image to bottom.

PIMO-positive mask for each histology slice was co-registered with the multiparametric maps of the corresponding MRI slice, according to a method described and successfully utilized before (*41*). Briefly, custom written MATLAB code was used to perform affine 2D registration based on manual detection of 4 corresponding landmarks in histology and MRI images. Prior to co-registration, slices where the tissues were broken or with missing parts were excluded from co-registration analyses.

DICE similarity coefficients (DSC) were calculated between each MRI slice and its corresponding histology slice by creating binary masks for both slices and measuring the similarity between the masks using DICE formula in equation 1. The similarity score ranges between 0 and 1. A score close or equal to 1 indicates the slices are very similar or identical. More specifically, if ***M*** is the binary mask of the MRI slice and ***H*** is the histology binary mask, the DSC score is obtained by the following DICE equation:

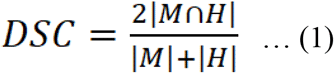

### 7. Hypoxic habitats using Deep Learning model

Deep residual network (ResNet), a type of CNN that uses residual blocks, achieves state-of-the-art performance in current image recognition field. In this study, the architecture of ResNet-18 with small number of filters in each layer was used to predict hypoxia probability of each pixel, which is shown in **Fig. S9**. In details, for each pixel within the tumor region, a 15×15 fixed size sliding window centered on this pixel was used to generate a multiparametric patch from T2* map, T2-weighted image, slope, time to max, and AUC from DCE-MRI images, which was fed into the DL model after z-score normalization for each channel to update the parameters with backward propagation. The binarized average value of the Pimonidazole-positive mask map within this window was encoded to one-hot and used as the label of this patch. The output of the network was used as the classification result to represent the hypoxia probability of each pixel. The final predicted hypoxic habitats could be reconstructed utilizing the location information of each pixel. To guarantee the accuracy of the labels, only samples with similarity score higher than the average scores were involved in the construction of the DL model.

During the training, binary cross entropy was employed as the loss function, while the Adam optimizer was used with an initial learning rate of 0.0001 and decaying by a factor of 0.2 if no improvement of the loss of the validation dataset was seen for 10 epochs. Additionally, augmentation including width/height-shift, horizontal/vertical-flip, rotation and zoom were used to expand the training dataset to improve the ability of the model to generalize. The implementation of this model used the Keras toolkit and Python 3.5.

### 8. Statistical analyses

Comparison of data between groups was done by using Student t-test or One-Way Analysis of Variance (ANOVA) followed by multiple comparisons test. Kaplan-Meier was used to estimate survival rates and the log rank test was used to analyze differences between the groups. P values <0.05 were considered statistically significant.

DSC was calculated to measure the spatial overlap between the predicted hypoxic habitats and PIMO-positive mask quantitatively. The cutoff to binarize the predicted hypoxia probability was determined according to the average optimal value to obtain the largest DSC for each training sample. The correlation between the true positive portion (i.e., PIMO-positive staining portion in histology) and the predicted positive portion by the model in the co-registered MRI was analyzed by Pearson correlation coefficient.

To measure the predictive ability of the predicted positive portion in identifying the samples with response to TH-302, area under the receiver operating characteristics curve (AUROC) was used. The optimal cutoff was determined to maximize the Youden’s index by balancing the sensitivity and specificity, and Cox proportional hazards model was used to analyze the prognostic value of the predicted positive portion.

## Supplementary Materials

**Fig. S1**. Body weight during therapy and comparison between day 0 and last day of therapy in the patient-derived xenograft (PDX) rhabdomyosarcoma model.

**Fig. S2**. Body weight during therapy and comparison between day 0 and last day of therapy in the radiation-induced fibrosarcoma cell line (RIF-1) model.

**Fig. S3**. Representative images showing CD-31 staining in PDX rhabdomyosarcoma and RIF-1 tumors.

**Fig. S4**. Tumor growth and survival plots for immunodeficient mice inoculated with RIF-1 cells.

**Fig. S5**. Correlation between true positive portion and predicted positive portion in the training, validation and test cohorts for (upper row) PDX tumors and (lower row) RIF-1 tumors.

**Fig. S6**. Quantification and representative PDX rhabdomyosarcoma tumors stained with DNA-damage marker (phospho γ-H2AX) and apoptosis marker Cleaved Caspase 3 (CC3).

**Fig. S7**. Individual thresholds were calculated based on Otsu method to identify pimonidazole-positive staining in each histology slice.

**Fig. S8**. Thresholding method (global or individual) did not affect calculation of pimonidazole-positive pixels.

**Fig. S9**. Schematic representation of the architecture of ResNet-18 used in this study.

**Table S1**. DICE similarity scores (DSC) calculated between each MRI slice and its corresponding histology slice for PDX tumor model.

**Table S2**. DICE similarity scores (DSC) calculated between each MRI slice and its corresponding histology slice for RIF-1 tumor model.

**Table S3**. Metrics for the CNN model for the PDX tumor model.

**Table S4**. Metrics for the CNN model for the RIF-1 tumor model.

## Acknowledgments

We thank Dr. Zaver M. Bhujwalla for providing the RIF-1 cell line, the Childhood Solid Tumor Network for providing the PDX cells and Molecular Templates for providing the drug evofosfamide. This work has been supported in part by the Small Animal Imaging Laboratory, **Image Response Assessment Team Core**, Analytic Microcopy Core, and Tissue Core Facilities at the H. Lee Moffitt Cancer Center & Research Institute, an NCI designated Comprehensive Cancer Center (P30-CA076292).

## Author contributions

B.V.J.P., W.M., D.R.R., G.V.M. and R.J.G. conceived and designed the experiments. B.V.J.P., W.M., S.H., W.D.V. developed the experiments. B.V.J.P., W.M., M.R.T., J.P., M.A.A., M.M.B., W.D.V., M.M.Bui, J.O.J., G.V.M. and R.J.G. analyzed and interpreted data. B.V.J.P., W.M. and R.J.G. wrote the manuscript with input from all authors. R.J.G. supervised the research.

## Data and materials availability

Data presented in this study are available from the corresponding author, upon request.

