## Supplemental Figures for "Targeting hypoxic habitats with hypoxia pro-drug evofosfamide in preclinical models of sarcoma"

**Supplementary Materials**

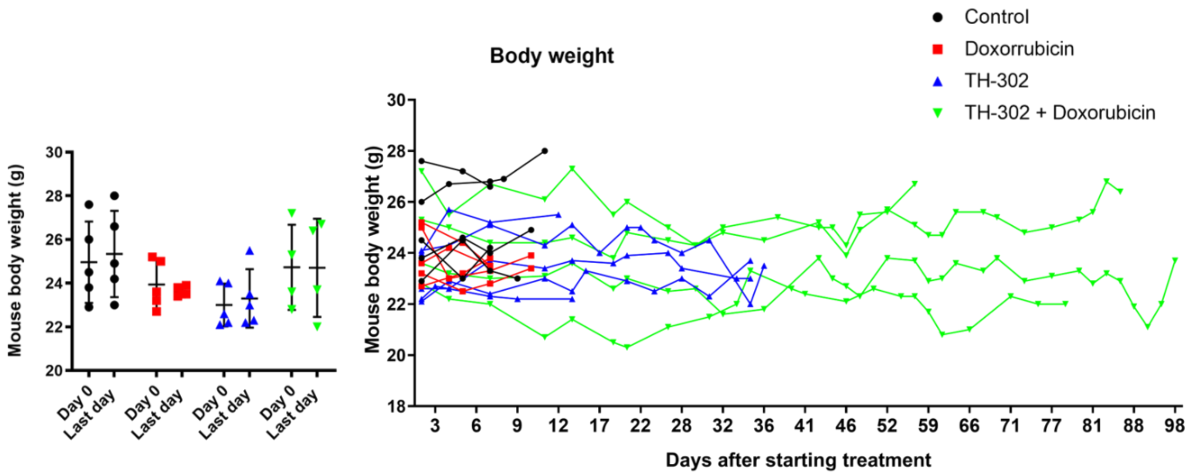


**Fig. S1.** Body weight during therapy and comparison between day 0 and last day of therapy in the patient-derived xenograft (PDX) rhabdomyosarcoma model. p-values of day 0 *vs* last day, p=0.76 Control; p=0.54 Dox; p=0.69 TH-302; p>0.98 TH-302 + Dox.

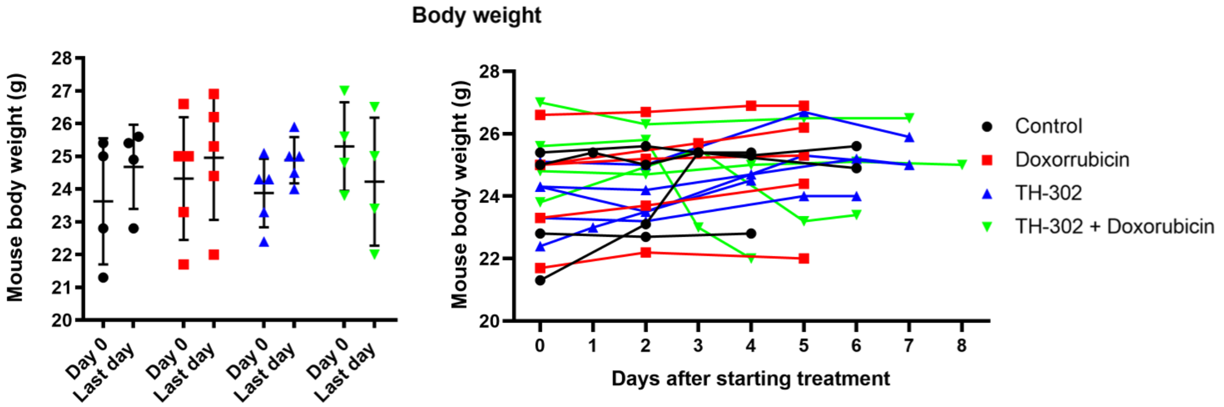


**Fig. S2.** Body weight during therapy and comparison between day 0 and last day of therapy in the radiation-induced fibrosarcoma cell line (RIF-1) model. p-values of day 0 *vs* last day, p=0.39 Control; p=0.60 Dox; p=0.11 TH-302; p>0.39 TH-302 + Dox.

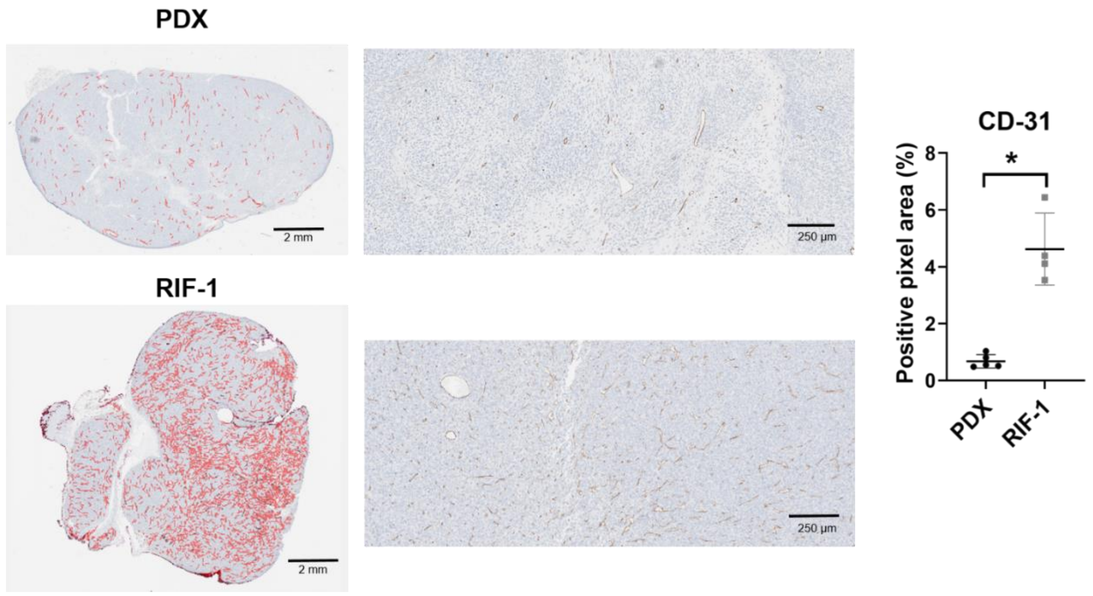


**Fig. S3.** Representative images showing CD-31 staining in PDX rhabdomyosarcoma and RIF-1 tumors. *p=0.002 by Student t test; Values of CD-31-positive area from control groups were compared between PDX and RIF-1 tumors in last day of therapy.

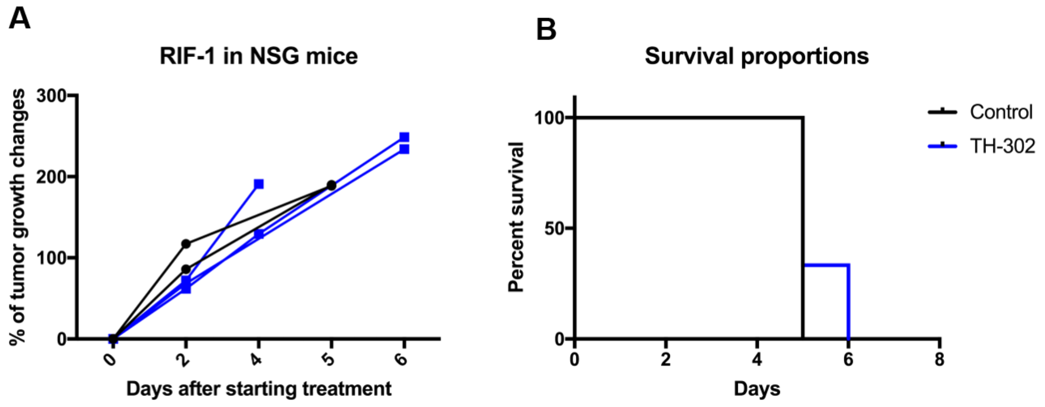


**Fig. S4.** Tumor growth and survival plots for immunodeficient mice inoculated with RIF-1 cells. **A.** Tumor growth changes (%) after starting treatment. No differences were observed between groups. **B.** Kaplan-Meier plots shown that there was not significantly difference in the OS between groups of treatment (p=0.41).

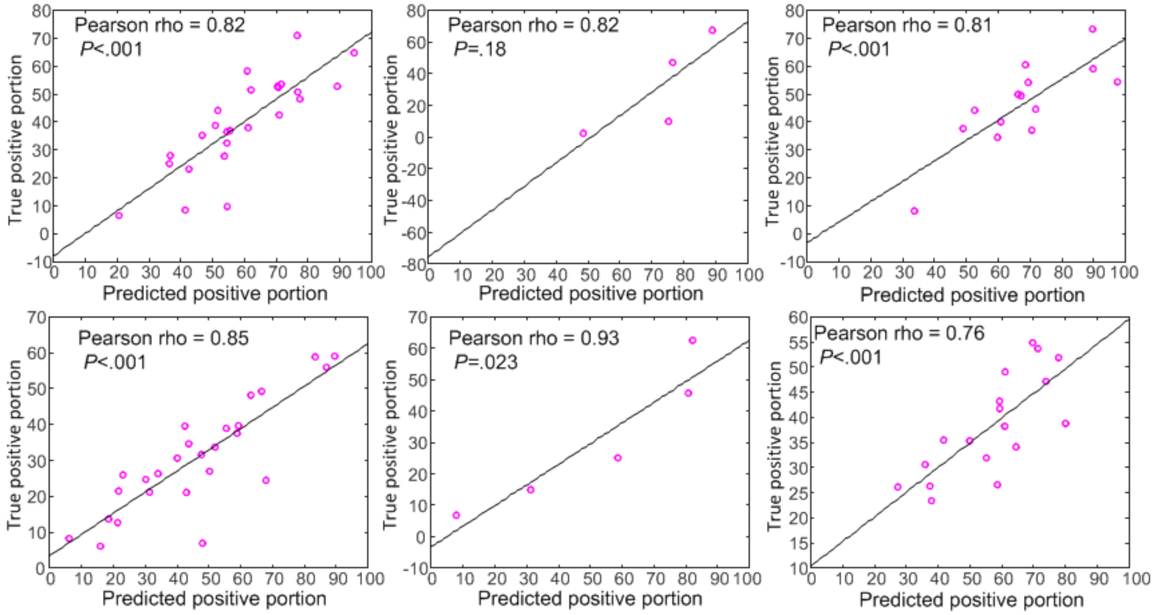


**Fig. S5.** Correlation between true positive portion and predicted positive portion in the training, validation and test cohorts for (upper row) PDX tumors and (lower row) RIF-1 tumors.

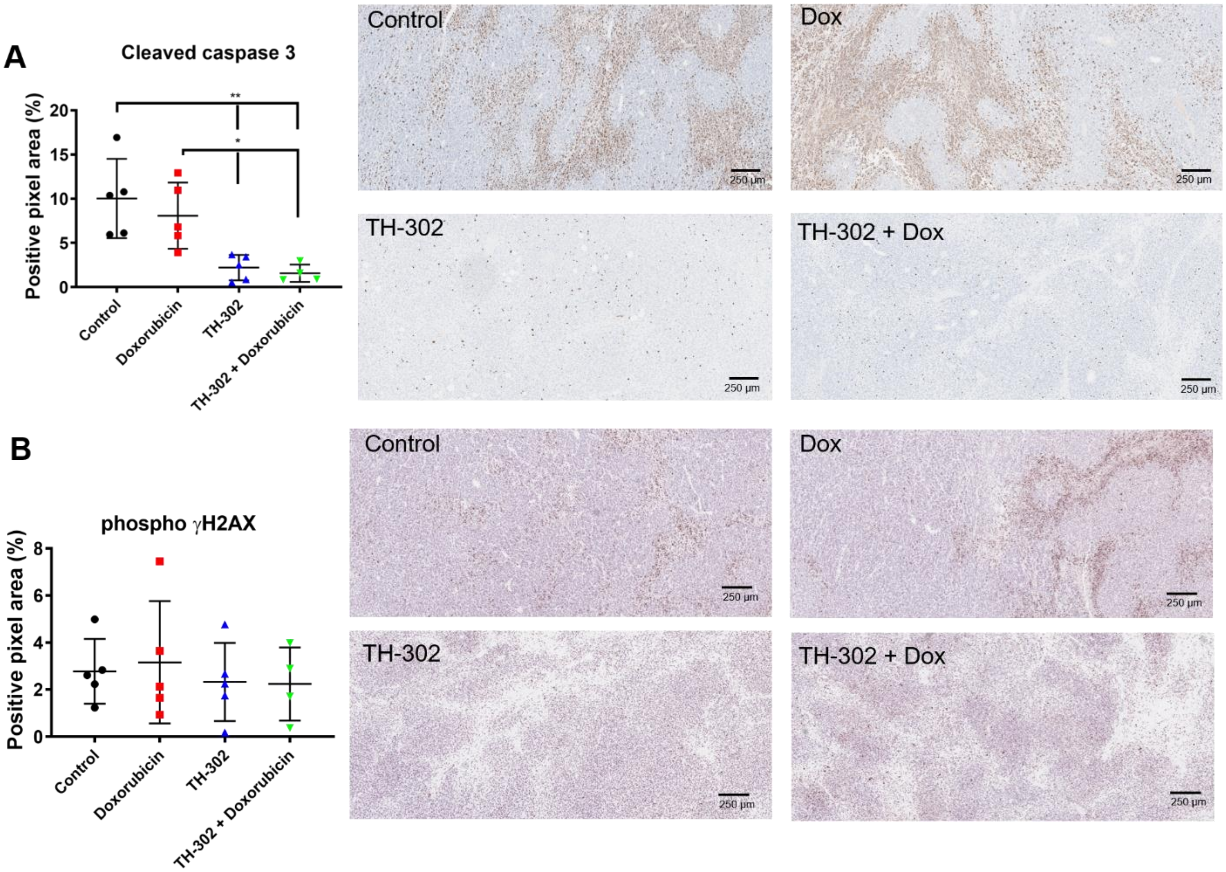


**Fig. S6.** Quantification and representative PDX rhabdomyosarcoma tumors stained with DNA-damage marker (phospho γ-H2AX) and apoptosis marker Cleaved Caspase 3 (CC3). **A.** CC3 comparisons (p>0.76 Dox *vs* Control; *p=0.006 TH-302 *vs* Control; *p=0.005 TH-302 + Dox *vs* Control; *p=0.04 TH-302 *vs* Dox; *p=0.03 TH-302 + Dox *vs* Dox; p=0.99 TH-302 *vs* TH-302 + Dox). **B.** Phospho γ-H2AX comparisons (p>0.98 Dox *vs* Control; p=0.98 TH-302 *vs* Control; p=0.97 TH-302 + Dox *vs* Control; p=0.89 TH-302 *vs* Dox; p=0.88 TH-302 + Dox *vs* Dox; p=0.99 TH-302 *vs* TH-302 + Dox). ANOVA followed by Bonferroni multiple comparison test. Values presented as mean± SD.

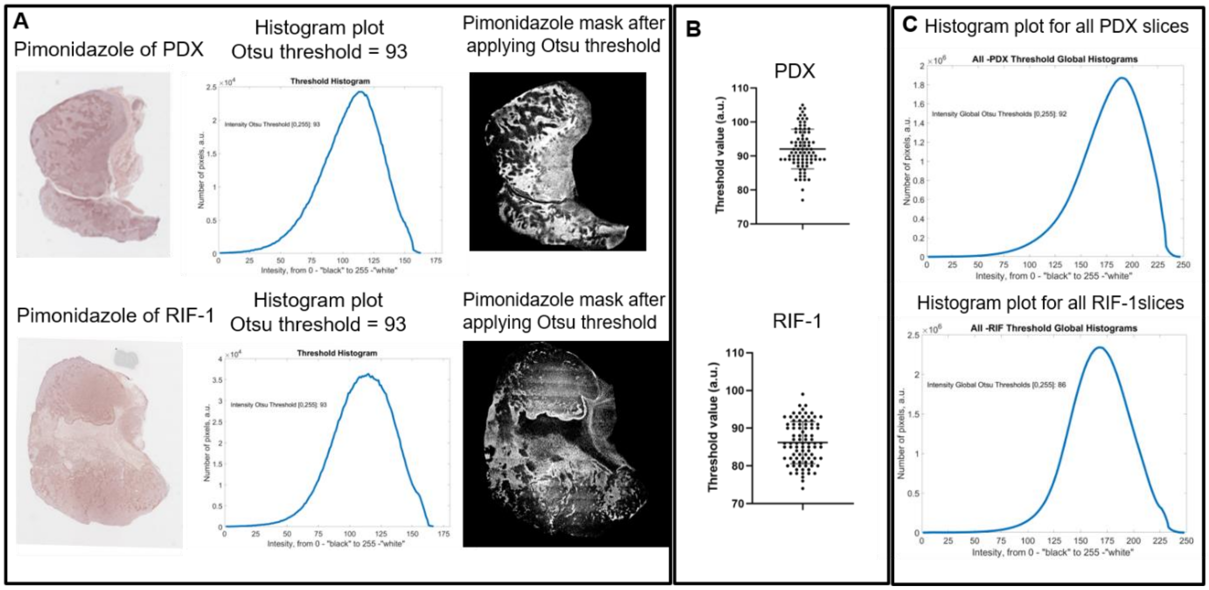


**Fig. S7. Individual thresholds were calculated based on Otsu method to identify pimonidazole-positive staining in each histology slice**. **A.** Representative histology slices stained with pimonidazole; histogram plots for individual histology slices to calculate threshold; and masked images with individual thresholds for PDX and RIF-1 tumors. **B.** Distribution of individual thresholds values for all histology slices for PDX and RIF-1 tumors. **C.** Histogram plots for all histology slices to calculate a global threshold for PDX and RIF-1 tumors.

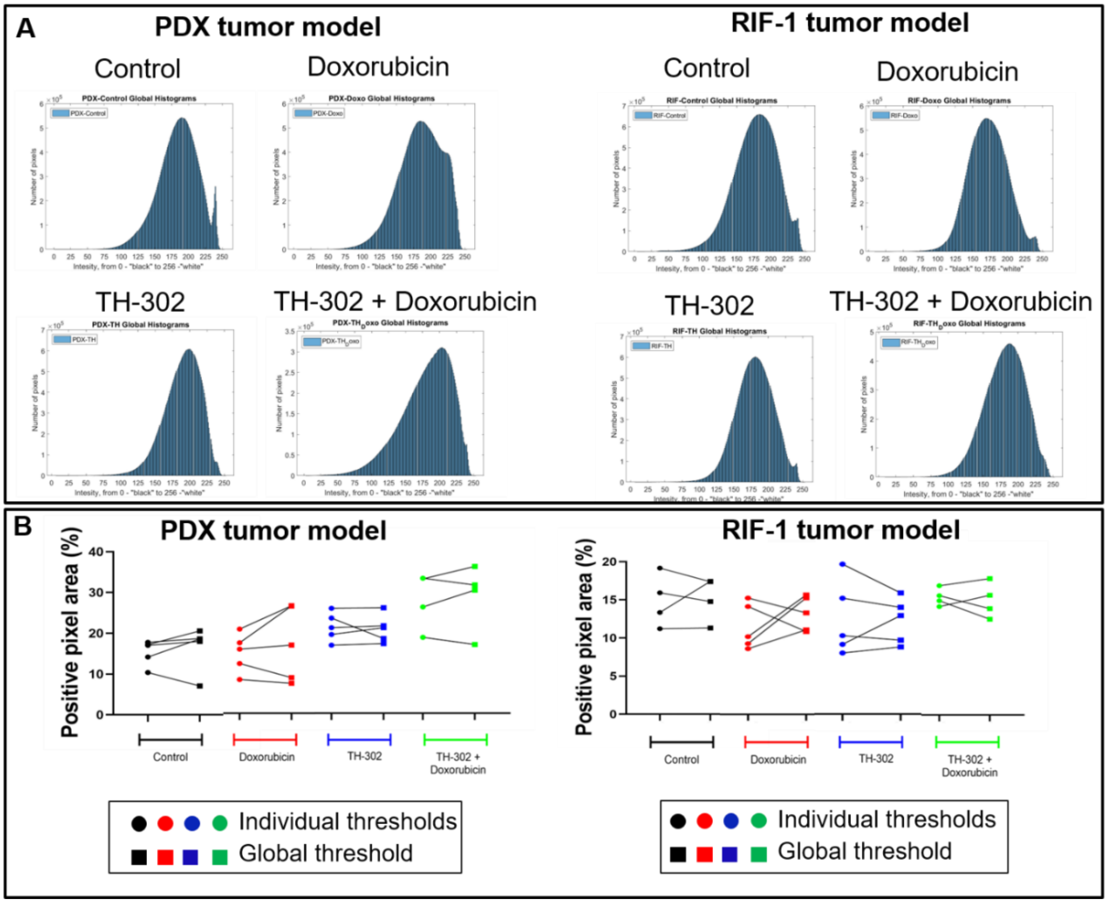


**Fig. S8. Thresholding method (global or individual) did not affect calculation of pimonidazole-positive pixels. A.** Histogram plots showing the distribution of pimonidazole pixels intensity for all histology slices in each group of therapy. **B.** Comparison of positive-pixel area (%) per groups when using individual thresholds for each slice to identify pimonidazole-positive areas (represented by circle) or a global threshold for all slices (represented by squares). There was not significantly difference in the percentage of pimonidazole-positive area in each group when using individual or global thresholding methods; p>0.05 by Paired t test. PDX tumor model (Control: p=0.38, mean of difference = 1.28; Dox: p=0.37, mean of difference = 2.27; TH-302: p=0.70, mean of difference = -0.47, TH-302 + Dox: p=0.58, mean of difference = 0.9), RIF-1 tumor model (Control: p=0.83, mean of difference = 0.30; Dox: p=0.41, mean of difference = 1.74; TH-302: p=0.87, mean of difference = -0.19, TH-302 + Dox: p=0.69, mean of difference = -0.42).

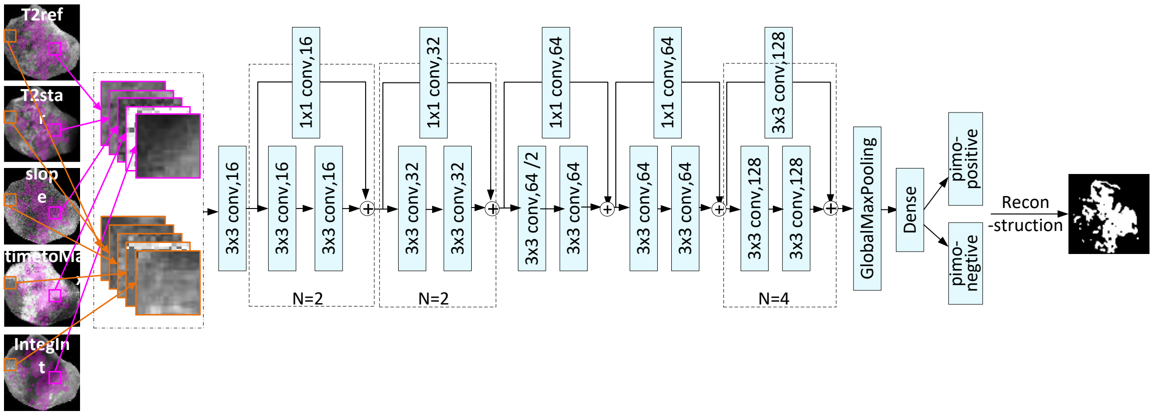


**Fig. S9.** Schematic representation of the architecture of ResNet-18 used in this study.
